# *De novo* design of Ras isoform selective binders

**DOI:** 10.1101/2024.08.29.610300

**Authors:** Jason Z. Zhang, Xinting Li, Alexa Rane Batingana, Caixuan Liu, Hanlun Jiang, Kevin Shannon, Benjamin J. Huang, Kejia Wu, David Baker

## Abstract

The proto-oncogene Ras which governs diverse intracellular pathways has four major isoforms (KRAS4A, KRAS4B, HRAS, and NRAS) with substantial sequence homology and similar *in vitro* biochemistry. There is considerable interest in investigating the roles of these independently as their association with different cancers vary, but there are few Ras isoform-specific binding reagents as the only significant sequence differences are in their disordered and highly charged C-termini which have been difficult to elicit antibodies against. To overcome this limitation, we use deep learning-based methods to *de novo* design Ras isoform-specific binders (RIBs) for all major Ras isoforms that specifically target the Ras C-terminus. The RIBs bind to their target Ras isoforms both *in vitro* and in cells with remarkable specificity, disrupting their membrane localization and inhibiting Ras activity, and should contribute to dissecting the distinct roles of Ras isoforms in biology and disease.

## Introduction

The Ras family of GTPases modulates the mitogen activated protein kinase (MAPK) and other intracellular signaling pathways essential for cell growth and survival, and mutations in Ras are prevalent in many human cancers. The four major Ras isoforms– KRAS4A, KRAS4B, HRAS, and NRAS–have high sequence homology and similar biochemical properties. Despite their similarities, the isoforms are differentially mutated in different cancers^1^, play different roles in drug resistance^2^, and have distinct subcellular locations^3,4^ due to their divergent disordered and highly charged C-termini. Whereas the structured portion of Ras GTPases has 90% sequence homology across the major Ras isoforms, the homology over the C-termini is only 8% (**Figure 1a-b**). While all isoforms can localize to the plasma membrane, NRAS, HRAS, and KRas4A are reversibly palmitoylated on their hypervariable C-termini, enabling endomembrane localization^3,4^. It has been challenging to develop Ras isoform specific reagents^5,6^; those that are available often result in multiple bands in immunoblotting experiments and are insufficient for more sensitive assays such as immunostaining. Despite decades of research into the different Ras isoforms, their specific signaling activities and functional roles remain unclear due to the lack of isoform-selective molecular tools, such as selective affinity reagents, complicating precise functional studies.

**Figure 1:**
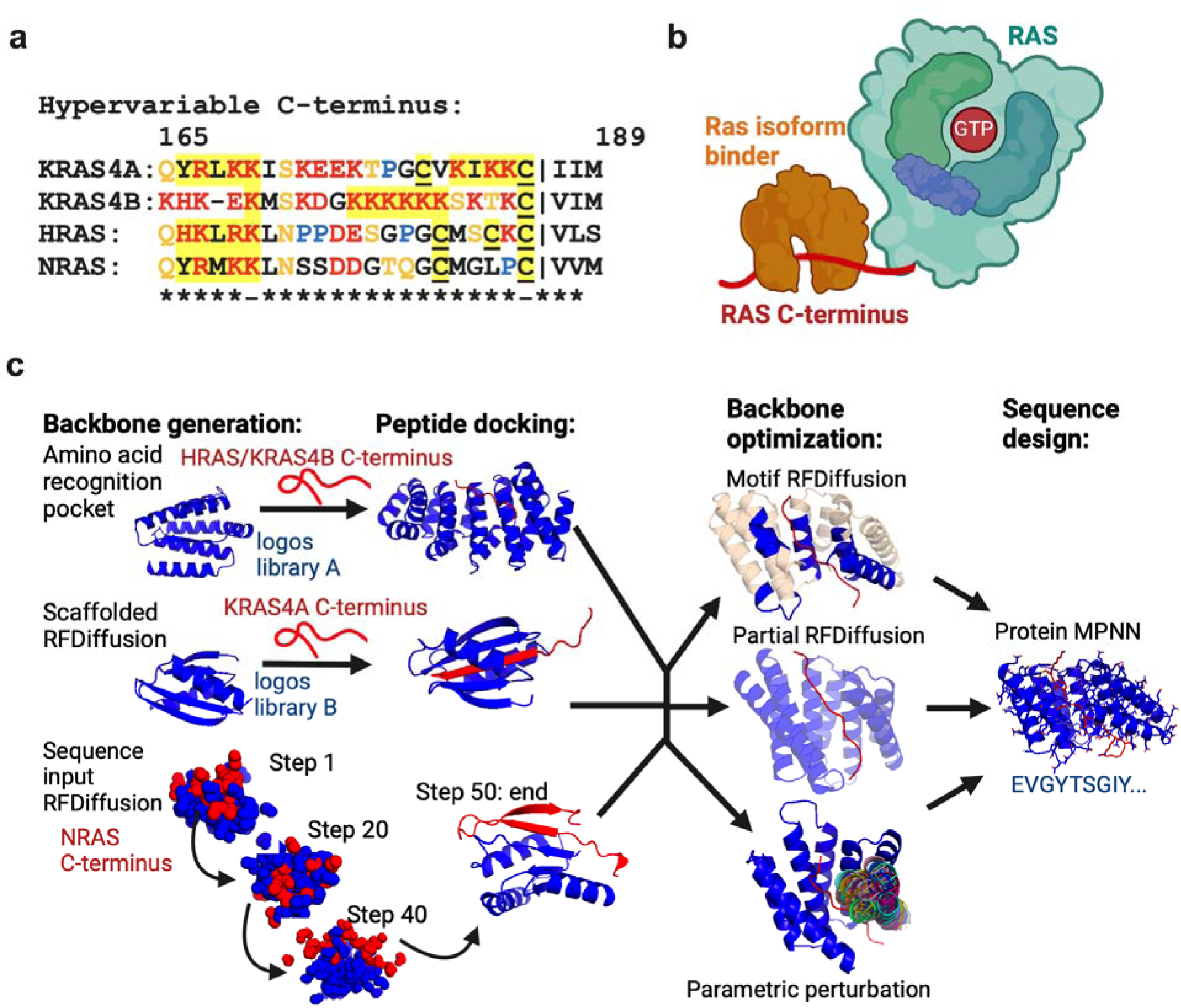
Computational design methods. **a)** Sequence alignment of the C-termini of the major Ras isoforms. Residues in red are charged, in orange are polar, and in blue are prolines. Highlighted in yellow are residues involved in membrane interaction. Underlined residues undergo lipid modification. **b)** Schematic representation of RIBs binding to the Ras C-terminus. **c)** Computational design. Backbone generation either through an amino acid recognition pocket based approach, scaffolded RFDiffusion, or sequence input diffusion. These peptide-scaffold backbones are then optimized for either through motif RFDiffusion (subsets of the backbone kept fixed), partial RFDiffusion (slightly altering the overall scaffold backbone), or parametric perturbation (rotating/moving a portion of the scaffold backbone in a specified manner). Next, ProteinMPNN was used to optimize the amino acid sequence for a given scaffold backbone. Scaffold-target complexes are evaluated using AlphaFold2 and Rosetta metrics to determine what to experimentally characterize.

Recent advances in protein design^7–9^ now enable the design of binders to a wide range of protein targets. Designing Ras isoform specific binders requires targeting the disordered and highly charged C-terminus of Ras (**Figure 1a**) as it is the only region that differs between the Ras isoforms. Targeting highly polar, native protein regions such as these C-termini of Ras (e.g. KRAS4B C-terminus is 86% polar residues, 63% charged residues) has been difficult^10^ due to the biochemical challenge of binder interaction competing with water interaction. Precise and extensive polar interactions are required for designing Ras binders, otherwise we need to pay substantial enthalpic penalty for striping away water-mediated hydrogen bonds and entropic penalty for stabilizing intrinsically disordered C-terminus of Ras. We reasoned that recently developed methods for designing binders to intrinsically disordered regions^11,12^ could enable design of Ras isoform specific binders superior to currently available antibodies. These reagents could be valuable tools for dissecting the roles of the different isoforms in cellular function and disease.

### Computational design of Ras isoform selective binders (RIBs)

We explored two different peptide binder design strategies for designing Ras isoform specific binders (RIBs). The first is “side-chain centered”, i.e., threads the sequences of disordered regions (IDRs) through ∼800 protein templates from logos^11^ (logos library A) with specific amino acid recognition pockets arranged to bind diverse sequences in a range of extended conformations. The second is “backbone centered”, i.e., generates backbones via either “scaffolded RFDiffusion” by threading the sequences into a subset of ∼200 β-sheet containing wrapping-up scaffolds also from logos^11^ (logos library B) or from random noise using sequence input RFdiffusion^12^ in which both the backbone conformation of the binder and the target were widely sampled. In both cases, the top scored designs would be optimized by the “refinement” protocol from the standard logos pipeline^11^, i.e., partial RFDiffusion^13^, motif RFDiffusion^11^, and parametric perturbation^14^. Sequences are all designed using ProteinMPNN^8^ (see Methods for details). Designs are selected for experimental characterization based on Alphfold 2 (AF2)^15^ prediction confidence metrics (pae interaction), predicted binding affinity (Rosetta *ΔΔ*G), and extent of buried polar atoms without hydrogen binding (buried unsaturated residues)^11^ (**Figure S1**). To achieve specificity, we prioritized binders with extensive polar interactions with the Ras C-terminus, but disregarding the last 4 residues which either get cleaved or are farnesylated.

Initially, we attempted to design binders to all the Ras isoforms using all methods. We found that both scaffolded and sequence input RFDiffusion methods primarily produced binder backbones that induced the target to conform to regular secondary structures.

This worked well with the KRAS4A and NRAS C-termini which were induced to form β-strands that formed an extended β-sheet with the binder, but not for the KRAS4B and HRAS C-termini which are less compatible with regular secondary structure (KRAS4B has a highly charged 6x lysine region and HRAS contains several unevenly spaced proline residues). The amino acid recognition pocket based approach was more successful in generating good scoring designs for all the Ras isoforms as it does not require any secondary structure propensity in the target. We selected for experimental characterization 8,317 (5,254 from sequence input and 3,063 from scaffolded) and 3,078 (343 from sequence input and 2,735 from scaffolded) designs made using RFDiffusion for KRAS4A and NRAS, and 2,556 and 1,264 designs made using the amino acid recognition pocket based approach for KRAS4B and HRAS.

### *In vitro* testing of *de novo* designed RIBs

We obtained synthetic DNA encoding the selected RIB designs and identified binders using yeast display^16,17^ (see Methods for details, **Figure S2a-b**). For KRAS4B and NRAS, the initial RIB library showed specific binding to the C-terminus but did not bind to full length Ras, perhaps due to steric interference with the full length proteins, which were not considered in our original design calculations. We carried out a second round of binder scaffold optimization via motif and partial RFDiffusion requiring compatibility with both the C-terminus and full length protein (**Figure S2c**), which resulted in smaller scaffolds clashing less with the target. For example, the helix of the KRAS4B C-terminus binder is predicted to clash with a portion of the structured domain of KRAS4B and hence portions of that helix were re-designed to accommodate binding for full length KRAS4B (**Figure S2d)**. Ultimately, yeast display selection yielded ∼250 binders for KRAS4A, ∼90 binders and NRAS, ∼50 binders for HRAS, and ∼20 binders for KRAS4B. For KRAS4A, binders were obtained from scaffolded RFDiffusion where a portion of the KRAS4A C-terminus adopts a β-strand conformation which forms extended β-sheet hydrogen bonds with the binder’s β-sheet (**Figure 2a and S3a**). In contrast, NRAS binders came from sequence input RFDiffusion and induced NRAS to adopt a conserved conformation where it formed multiple β-strands to create extended β-sheets with the binder (**Figure 2a and S3a**). The successful KRAS4B and HRAS binders came from the amino acid recognition pocket approach and induced KRAS4B or HRAS to adopt a non-regular secondary structure with an extended interface (**Figure 2a and S3a**).

**Figure 2:**
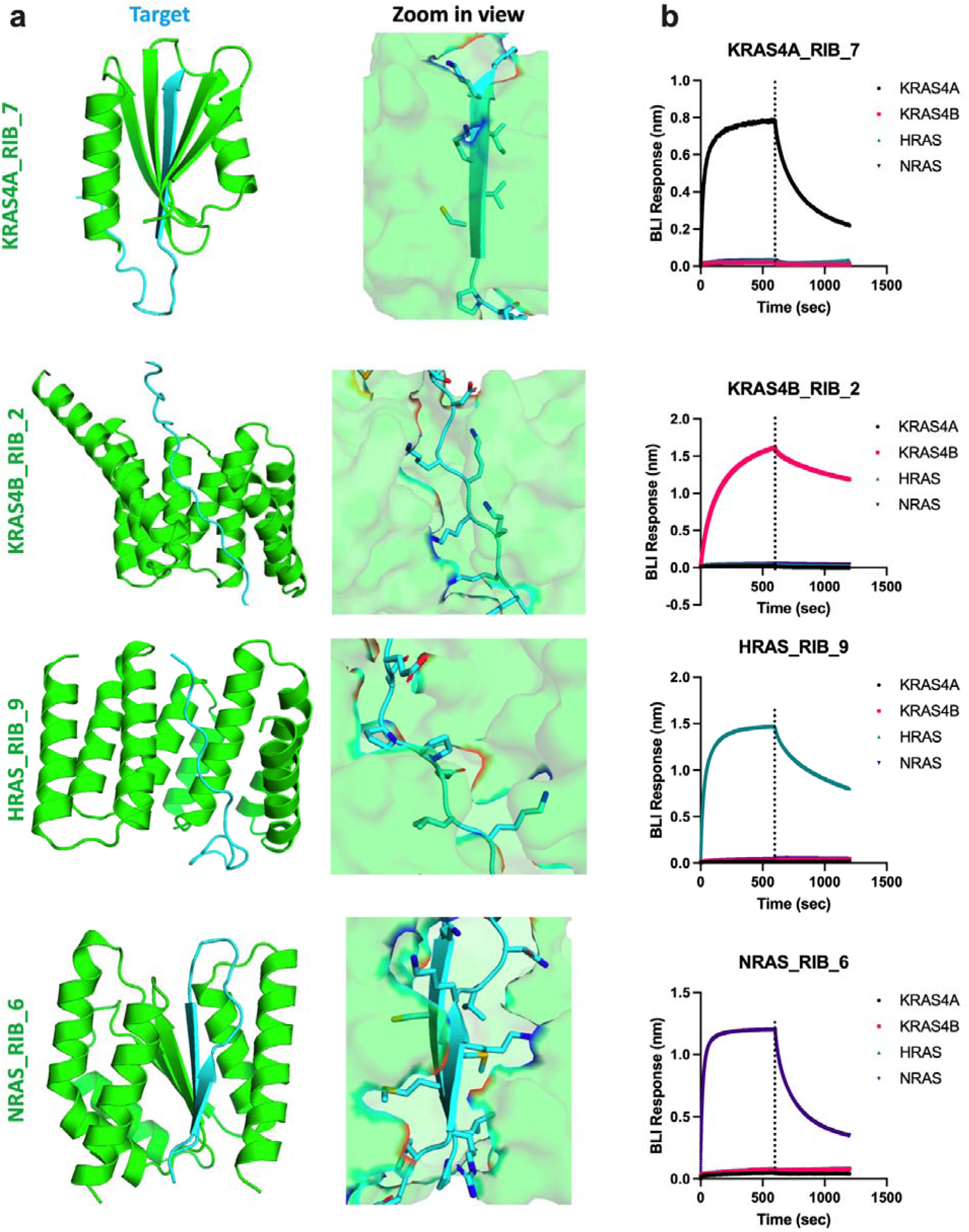
RIBs specifically bind to their target *in vitro*. **a)** Design models of the RIBs (green) in complex with their Ras C-termini targets (cyan). **b)** Representative biolayer interferometry (BLI) results of RIBs (1µM) interacting with the different Ras isoforms full length protein. In each case, binding is specific for the intended target. Each experiment was repeated 3 times.

We next expressed and purified the top 10 RIB hits from yeast display per target for *in vitro* characterization. Corroborating the yeast display results, RIBs specifically bound to their cognate full length target (**Figure 2b**) with affinities ranging from 35-1300nM (**Figure S3b-c, Table S1**) measured using bilayer interferometry. The RIBs bound to their intended target in a nucleotide-independent manner, which is expected as the C-terminus of Ras is distal from the nucleotide binding site^1^ in the structured portion (**Figure S3d**). All 10 RIBs (numbered based on BLI-derived kd ranking) for each target that were tested experimentally bound both *in vitro* and in cell experiments (**Table S1**), and the RIBs with the most consistent and specific binding throughout the experiments are shown in **Figure 2** and are used for the rest of the paper (**Figures 3-4 and S5-7**). Of note, the binders with tight affinity (kd<100nM, **Table S1**) were not necessarily the most specific when applied in cell experiments. In the design model of the scaffolded RFDiffusion generated KRAS4A_RIB_7:KRAS4A complex, the C-terminal section of the KRAS4A C-terminus forms a β-strand that pairs with β-strands deep within the binder and is packed by surrounding helices. In the design model of the amino acid recognition pocket-derived KRAS4B_RIB_2:KRAS4B and HRAS_RIB_9:HRAS complex, a portion of the KRAS4B or HRAS C-terminus forms an extended unstructured interface with the binder. In the design model of the sequence input RFDiffusion-generated NRAS_RIB_6:NRAS complex, the NRAS C-terminus forms two β-strands (innermost β-strand is NRAS C-terminus) which is incorporated into an extended β-sheet with the binder and is packed by surrounding helices.

**Figure 3:**
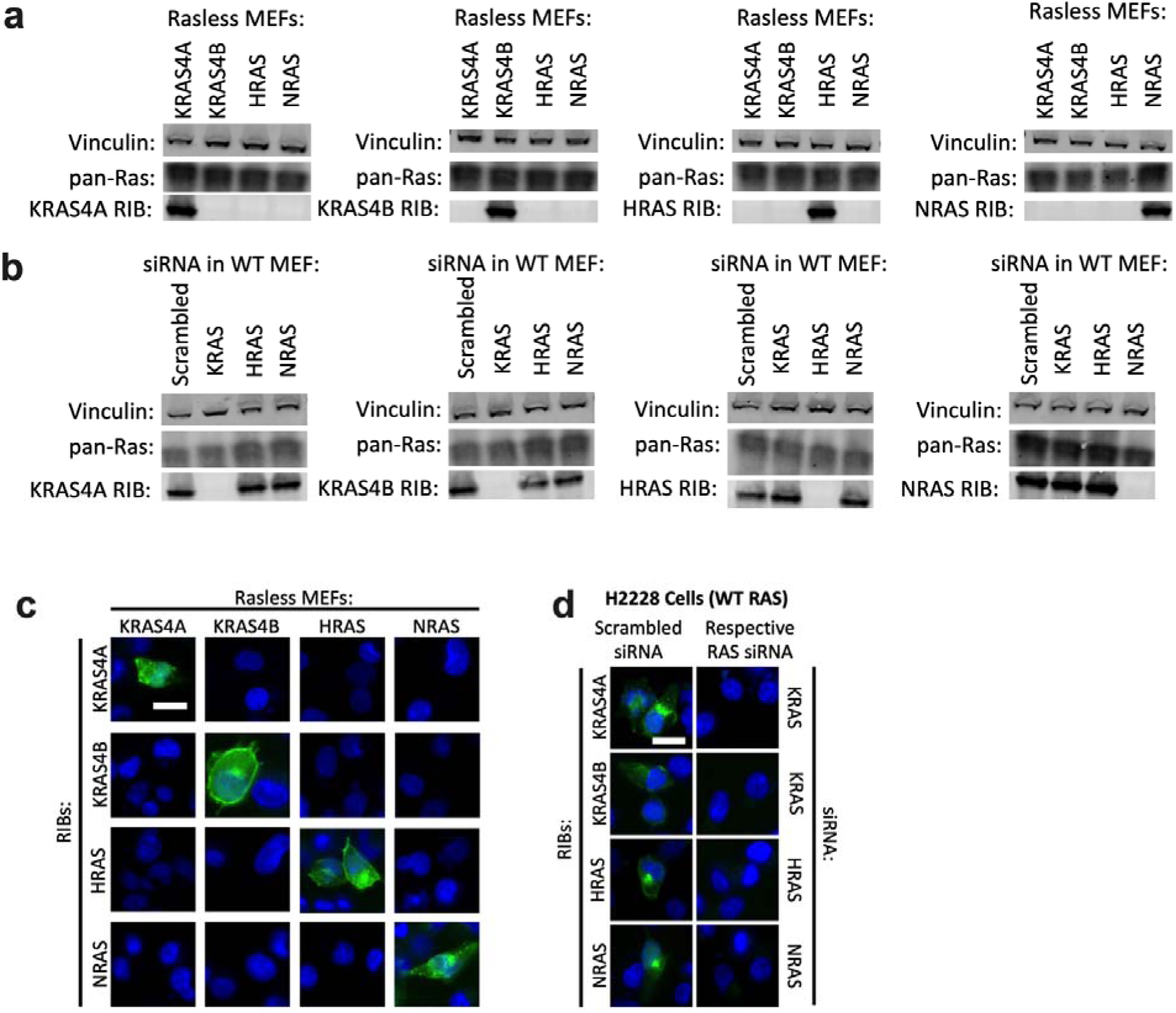
RIBs specifically bind to their targets in cells. **a)** Lysates from Rasless MEFs, which express only one WT Ras isoform at a time, were run on SDS-PAGE gels and the Ras isoforms were probed with 10µM biotinylated RIBs or antibodies. Shown are representative blots from 3 independent experiments. **b)** Wildtype (WT) MEF cells were transfected with either Scrambled or Ras isoform specific siRNA. Two days later, cells were lysed, ran on SDS-PAGE gels, and probed with biotinylated RIBs or other antibodies. Shown are representative blots from 3 independent experiments. **c)** Rasless MEFs were probed via biotinylated RIBs (green) and DAPI (blue). Shown are representative epifluorescence images from 3 independent experiments. Scale bar=10µm. **d)** Wildtype (WT) H2228 cells were transfected with either Scrambled or Ras isoform specific siRNA. Two days later, cells were fixed, permeabilized, and probed with biotinylated RIBs (green) and DAPI (blue). Shown are representative epifluorescence images from 3 independent experiments. Scale bar=10µm.

**Figure 4:**
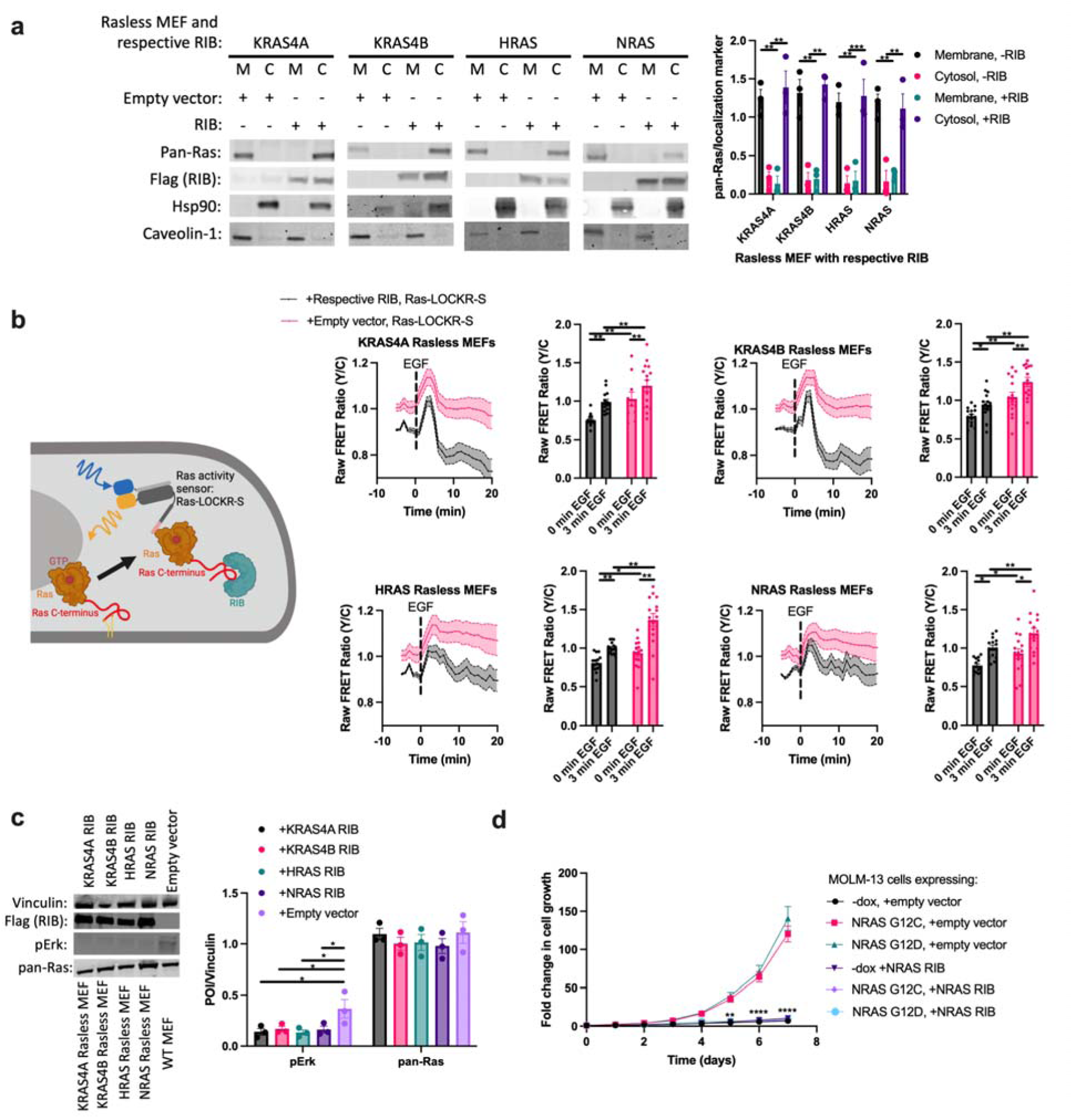
RIB expression alters Ras localization and signaling. **a)** Rasless MEFs were transfected with either empty vector (-RIB) or their respective RIB (+RIB) for 1 day, lysed, and underwent membrane fractionation and immunoblotting analysis. Left: representative immunoblot. Right: densitometry quantification of immunoblots comparing pan-Ras signal with the signal from the localization marker (Hsp90 for cytosol, Caveolin-1 for membrane) (n=3 experiments). Statistics: two-way ordinary ANOVA. **b)** Rasless MEFs were transfected with Ras-LOCKR-S and either RIB fused to mCherry or empty vector. Then cells were imaged and stimulated with 100ng/mL Epidermal Growth Factor (EGF). Top left: Schematic of experiment. Left: Time-course imaging of raw FRET ratios (yellow over cyan (Y/C)) representative of three independent experiments. Right: Bar graph quantification of the Ras-LOCKR-S raw FRET ratios across all experiments (n=15 cells per experiment). Statistics: two-way ordinary ANOVA. **c)** Rasless MEFs were transfected with their respective RIB for 1 day. These cells were compared to WT MEFs. These cells then underwent immunoblotting analysis. Right: densitometry quantification of immunoblots (n=3 experiments). Statistics: two-way ordinary ANOVA. **d)** MOLM-13 cell lines expressing either NRAS G12C or G12D were treated with or without doxycycline to induce mutant NRAS expression. Cells were also transfected with NRAS RIB expressing plasmid or empty vector and co-treated with AC220 (quizartinib, FLT3 inhibitor)^29,30^ (see Methods for details). Cells were then counted over a 7 day period (n=3 experiments). Statistics: one-way ordinary ANOVA.

### RIBs identify RAS isoforms in mammalian cells

To test the ability of RIBs to selectively bind Ras isoforms in mammalian cells, we used mouse embryonic fibroblast cell lines (MEFs) in which only one of the major wildtype (WT) Ras^18,19^ isoforms is expressed at a time (e.g. KRAS4A Rasless MEFs express only WT KRAS4A). To test for RIB specificity, we carried out the analogue of an immunoblot in which Rasless MEF cell lysates separated on SDS gels were probed with biotinylated RIBs (see Methods for details) followed by labeling by AlexaFluor 488-conjugated streptavidin. For each RIB, little binding was observed for the non-cognate cell lines, and a single band with the molecular weight of Ras was observed in the cognate cell line (**Figure 3a**). To determine whether RIBs can detect endogenous levels of the target, we carried out siRNA knockdown experiments (**Figure 3b**) in unmodified WT MEFs which contain all the major Ras isoforms. RIB-mediated band signal was lost only when its target was knocked down (e.g. only NRAS siRNA eliminates NRAS RIB-mediated band). To test whether RIBs can detect endogenous levels of Ras isoforms, we probed cell lines with different Ras genetic backgrounds and observed only one major band was seen around 20 kDa, the expected molecular weight of Ras (21 kDa) (**Figure S4**). In contrast, the most specific known Ras isoform antibodies^5^ displayed many non-specific bands across the same cell lines (**Figure S4**), highlighting the specificity of RIBs compared to other reagents.

### Imaging Ras isoforms in cells

We investigated whether the RIBs are sufficiently specific and sensitive enough for fluorescence imaging assays. We applied biotinylated RIBs to fixed and permeabilized Rasless MEFs and used AlexaFluor 488-conjugated streptavidin for imaging detection. The RIBs only stain their respective Rasless MEFs (e.g. KRAS4A RIB fluorescence signal is only seen in KRAS4A Rasless MEFs) (**Figure 3c**), with no significant signal in the other Rasless MEFs (**Figure S5a**). As found in other studies^5^, only a subset of cells showed significant staining signal possibly due to heterogeneity in Ras isoform expression. The specific fluorescence labeling with the RIBs is greater than that of the previously described isoform-specific antibodies^5^ which do not show significant immunostaining signal in the corresponding Rasless MEF cell line (**Figure S5b**). Only the NRAS antibody showed significant immunostaining signal, but knockdown of NRAS via siRNA did not diminish immunostaining signal, suggesting lack of specificity (**Figure S5b**).

The fluorescence staining results suggested differences in localization with KRAS4B having the most plasma membrane localization and NRAS the most endomembrane signal, consistent with previous work^2,3^. However, localization using the RIBs could be biased if RIB binding is reduced by Ras C-terminal palmitoylation since we did not include covalent modification by palmitoylation in our initial design computations and non-palmitoylated Ras is thought to be primarily in the golgi and ER while palmitoylated Ras is in the plasma membrane^3^. To test for this potential bias, we probed Rasless MEFs both with our biotinylated RIBs and a pan-Ras antibody, which binds to the structured domain of Ras thus should not interfere with RIB binding and should only label the expressed Ras isoform in these Rasless MEFs (**Figure S6a**). The signal from KRAS4A, KRAS4B, and HRAS RIBs mostly co-localized with the signal from pan-Ras immunostaining, suggesting that the RIBs do not have a significant bias in subcellular binding location. The NRAS RIB led to more endomembrane signal and less plasma membrane signal than pan-Ras immunostaining. The predicted complex structures of NRAS RIB with NRAS C-terminus shows that the NRAS palmitoylation site C181^3,20,21^ is buried within the interface, thus we hypothesized that the lower signal colocalization could be due to the NRAS RIBs binding primarily to the non-palmitoylated version of NRAS, which should be more endomembrane localized. In BRAFV600E-expressing Rasless MEFs, cells which do not express any of the major Ras isoforms, either ectopically expressing NRAS WT or palmitoylation-resistant C181S, NRAS RIBs pulled down NRAS C181S^3,20,21^ to a greater extent than NRAS WT (**Figure S6b**), suggesting indeed that NRAS RIBs preferentially bind to the non-palmitoylated forms of NRAS.

To assess whether our RIBs can label in intact cells endogenous levels of Ras isoforms, H2228 cells which contain all major Ras isoforms were transfected with Ras isoform selective siRNAs. While each of the RIBs had detectable staining signal when scrambled siRNA was transfected, knockdown of the targeted Ras isoform in H2228 cells eliminated this signal (**Figure 3d**), demonstrating that our RIBs can measure endogenous Ras isoforms. This has not been achievable with previously described isoform-specific antibodies^5^.

### Overexpression of RIBs disrupts Ras localization and signaling

We next explored the effects of ectopically expressing the RIBs inside cells. As the RIBs bind to the Ras C-terminus which is important for membrane localization, these RIBs could disrupt their membrane localization. Rasless MEFs expressing their respective RIBs shifts the Ras isoform from being in the membrane to the cytosolic fraction (**Figure 4a**). Furthermore, Rasless MEFs expressing GFP tagged RIB (RIB-GFP) and immunostained with pan-Ras antibodies, which report on the localization of the only Ras isoform expressed, showed a significant shift in Ras localization from membranes to the cytosol for all isoforms (**Figure S7a**). Quantitative membrane co-localization analysis confirmed that membrane localization of Ras was statistically significantly decreased when RIBs were expressed (**Figure S7b**). These results suggest that RIB overexpression disrupts Ras membrane localization.

The canonical view is that Ras requires a membrane to be in its active GTP loaded state^1,22,23^, but a membrane-less cytosolic pool of Ras^21,24^ could also be active. As RIB expression enhances the cytosolic localization of Ras, we explored whether this perturbation in Ras localization affects signaling. We used a previously constructed Ras activity biosensor (Ras-LOCKR-S^24^) to measure Ras-GTP loading in live cells with single cell resolution. Rasless MEFs expressing either mCherry tagged RIB or empty vector and co-expressing untargeted Ras-LOCKR-S to measure whole cell Ras activity were imaged (**Figure 4b**). For all the Ras isoforms tested, the Ras-LOCKR-S FRET ratios (yellow over cyan (Y/C)), which reports on Ras-GTP loading, were lower in cells overexpressing RIB (“0 min EGF” in **Figure 4b**). However, upstream Ras activation by 100ng/mL epidermal growth factor (EGF) still induced some transient Ras activation (“3 min EGF” in **Figure 4b**) when RIBs were expressed, and this is true for all Ras isoforms tested. While KRAS4A and KRAS4B Rasless MEFs displayed transient activation of Ras with Ras signaling going to or even below baseline Ras activity, NRAS and HRAS Rasless MEFs showed transient Ras activation where Ras activity still lingers 10 minutes after EGF stimulation^25^ (**Figure 4b**). These differences in Ras activity dynamics may be due to the differential localization of these Ras isoforms. A negative control (NC) version of Ras-LOCKR-S^24^ that is a point mutant away from Ras-LOCKR-S but renders the biosensor insensitive to Ras-GTP displayed no difference in FRET ratios between RIB versus empty vector expression (**Figure S7a**).

Looking downstream of Ras activity, we probed for Erk activity changes during RIB expression by probing for phospho-Erk (pErk) levels. Rasless MEFs expressing their respective RIB displayed decreased pErk/Vinculin levels compared to WT MEFs transfected with empty vector (**Figure 4c**). However, EGF stimulation in these same conditions led to increases in pErk/Vinculin levels that were indistinguishable from WT MEFs transfected with empty vector (**Figure S7d**), corroborating our Ras-LOCKR-S data showing that Ras activity can still be stimulated when RIBs are expressed. This Erk activation observed when expressing RIBs is dependent on Ras farnesylation as dual farnesyl transferase and geranylgeranyl transferase-1 inhibition greatly diminishes pErk levels (**Figure S7e**), aligning with previous reports indicating Ras activation and downstream signaling requires Ras farnesylation^26^ (the last 3 or 4 C-terminal residues which get farnesylated and cleaved were not included in RIB design, thus we expect that RIB binding to Ras isoforms is not influenced by the farnesylation). Probing with a pan-Ras antibody showed no significant differences in Ras levels (**Figure 4c and S7d**), suggesting that these decreases in basal pErk/Vinculin levels is not due to Ras protein expression differences. To verify that RIB overexpression only affects their respective Ras isoform, RIBs were overexpressed in all the Rasless MEFs and probed for Ras activity levels via Ras-LOCKR-S or Erk activation via pErk/Erk levels (**Figure S7f**). Only RIB expression in their respective Rasless MEF decreased Ras activity and Erk activation, demonstrating the specificity of the designs. Overall, these results corroborate the dogma that Ras-GTP loading is enhanced when in proximity to membranes, but also suggest that cytosolic Ras can be activated and lead to downstream signaling.

Membrane interaction of Ras is important for its activation and downstream signaling, which ultimately alters cell growth and proliferation. As RIBs can disrupt Ras localization and signaling, we wondered if RIB expression can alter cell proliferation. We focused on NRAS as our NRAS RIBs have a clear preference for non-palmitoylated NRAS (**Figure S6B**) and NRAS depalmitoylation has emerged as a promising therapeutic strategy for melanoma, hematologic cancers, and other *NRAS*-mutant cancers^27,28^. We expressed NRAS RIB in engineered isogenic MOLM-13 cell lines that are dependent on doxycycline-induced expression of mutant NRAS or KRAS4B. We found that NRAS RIB selectively inhibited the growth of mutant NRAS expressing MOLM-13 cell lines (**Figure 4d**) but not KRAS4B expressing MOLM-13 cell lines nor MOLM-13 cells not given doxycycline (**Figure 4d and S8**). These results indicate that perturbing the localization of Ras can influence Ras activation, downstream signaling, and cell growth.

## Discussion

The Ras isoforms exhibit distinct subcellular distributions and mutational preferences in cancer yet tools to differentiate these Ras isoforms have been lacking due to their high sequence similarity except at their C-terminus. The *de novo* designed RIBs bind to these highly charged and disordered C-termini with higher specificity than any reagents described to date^5,6^ both *in vitro* and in cells. Expression of the RIBs enhanced the cytosolic localization of these Ras isoforms which led to decreased Ras activity, corroborating the notion that membrane association of Ras is important for its function. Given the central importance of Ras in signaling and disease, our Ras isoform selective binders should be useful for a wide range of applications including isoform specific inhibitors (**Figure 4b-c**), affinity handles for targeted degradation, diagnostic markers for cancer patient samples, and Ras isoform specific biosensors.

## Acknowledgements

We acknowledge funding from HHMI (J.Z.Z. and D.B.), Helen Hay Whitney Foundation (J.Z.Z.), NCI K99-CA293001 (J.Z.Z.), the Croucher Fellowship (H.J.), the Audacious Project at the Institute for Protein Design (J.Z.Z, D.B.), NIH K08-CA256489 (B.J.H.), and NIH R01 CA193994 (K.S., B.J.H.). We thank D.J. Maly, F. McCormick, and D. Esposito for fruitful discussion of the Ras results, C.M. Dobbins and S. Cheng for help with mammalian cells and cell culture, and the RAS Initiative at the Frederick National Lab for providing the MEF cells.

## Author contributions

J.Z.Z. conceived of the project. K.W. supervised design strategies for making Ras isoform specific binders in this work, designed and assembled templates for KRAS4B, designed the scaffolds used in the logos pipeline. J.Z.Z. computationally designed these proteins with help from K.W., H.J. and C.L.. J.Z.Z. and D.B. supervised, designed, and interpreted the experiments. J.Z.Z. performed all experiments. X.L. made the Ras isoform peptides. J.Z.Z., and D.B. wrote the original draft. All authors reviewed and commented on the manuscript.

## Methods

### Amino acid recognition pocket based design and scaffolded RFDiffusion

As described in the previous work featuring the logos pipeline^10,11^, the recognition pocket approach took advantage of the overall flexible nature of disordered regions as well as the base of amino acid recognition in a string of polymers. In short, this approach would thread the query target into a diverse and robust family of ∼1,000 de novo protein-peptide complex templates, made up of arbitrary combos of 20 amino acid recognition pockets assembled into computationally designed universal binding modes. Initial docks were judged by how well fit each individual amino acid from the query target was, and how many of those fits were out of non-fits. Top docks were then identified by the pipeline, which were considered as the best design starting points. Sequence design, structural prediction, RFdiffusion refinement were then applied sequentially, until new customizable binding proteins were made and predicted well to the query target of interest.

#### Sequence Design

All accepted docks are sent to ProteinMPNN with a customized script that applies empirical weights to certain amino acids. Two iterations of ProteinMPNN and FastRelax are performed to adjust chain B backbones further and optimize the binder SAP score. Sequences with the lowest ProteinMPNN scores are filtered and sent to our customized AF2 package. This filtering process varies depending on the target’s difficulty level. Criteria can be adjusted based on individual targets and design campaigns.

#### Predictions

After two cycles of sequence design and AF2 (prediction-guided sequence design), passing designs undergo AlphaFold-multimer for final predictions. For highly polar or proline-rich targets, AlphaFold-multimer proved more predictive than AlphaFold-initial guess. AlphaFold-monomer is then applied to assess designed monomer predictions and select the final designs for experimental testing.

#### Scaffolded RFDiffusion

A subset of ∼200 β-sheet containing templates derived from the amino acid recognition pocket based design of template backbones (logos library B) were used as the starting point. These templates then went through the same pipeline as described above of peptide threading, sequence design, predictions, and then diffusion refinement.

### KRAS4B binder design

Following the same computation methodology as the original logos pipeline, manual refinement was performed on designs against this target. Due to the highly charged nature of this target (fill in the AA), we reasoned that a set of customized assembly of positive charged pockets (which was unenriched in the original logos library) would boost design success rate. Therefore, all the mono-, di-peptide pockets containing target of positively charged amino acids (i.e., K, R, H) were identified, same pocket assembly strategy as in previous work was applied^10,11^, along with another set of logos templates toward targets containing >60% of positively charged amino acids. For these two sets, chain B (target) was replaced with the KRAS4B target. The same diffusion refinement (i.e., motif diffusion for 8,000 trajectories each template, one-sided partial diffusion for 1,000 trajectories each template) was performed followed by ProteinMPNN and Alphafold2.

#### Sequence input RFDiffusion based design

For each target, approximately 10,000-50,000 diffused designs were generated based solely on the sequence input of the target. The resulting backbone library was subjected to sequence design using ProteinMPNN, followed by AF2 initial guess. These designs were then filtered based on interface pAE and pLDDT. Additionally, AF2 monomer analysis was conducted using only the binder sequence without the peptide, allowing for filtering based on the binder’s monomer pLDDT and RMSD relative to the binder design model. Subsequently, FastRelax was employed to obtain Rosetta metrics. The resulting binders were further filtered according to specific criteria, including contact molecular surface, ddG, SAP score, and the number of hydrogen bonds. These filtering criteria were meticulously chosen to narrow the selection down.

#### Backbone optimization with RFDiffusion and parametric perturbation

The process of pocket recognition and assembly played a crucial role in refining precise interactions with individual amino acids in a sequence-specific manner. This was particularly important when dealing with challenging polar or charged targets or targets that would not form regular secondary structures upon binding.

#### Motif RFDiffusion

As described in the previous work of *logos* pipeline^10,11^, if the design did not meet the complex prediction criteria mentioned above, or if a larger number of tested designs was preferred, motif RFDiffusion was employed to optimize the fit between less-than-ideal interacting pockets and binder backbones. In this work, especially for the need of yeast display screening thousands of binders a time, we used motif RFDiffusion more dramatically than originally developed to meanwhile shrink the binder size. Here, we retained a significant portion of the original binding interactions and motifs mostly around the central part of the RIBs to preserve the advantages of our general platform. Meanwhile, we deleted the terminal region and let RFDiffusion regrow with a smaller length. As part of this strategy, we developed partial RFDiffusion with small steps and multi-motif-constrained RFDiffusion (motif RFDiffusion).

#### Partial RFDiffusion (One-Sided)

As in other cases, RFDiffusion was modified to allow the input structure to be noised only up to a user-specified time step, rather than completing the full noising schedule. Consequently, the denoising trajectory’s starting point retained information about the input distribution, leading to denoised structures that remained structurally similar to the original input. In our work, the time step was set lower to maintain structural similarity, given that the high design resolution of hydrogen bonding interactions (especially with polar amino acids) was deemed critical. Specifically, 10-18 noising timesteps (10, 12, 15, 18) out of a total of 50 were selected for this paper’s work. For each target parameter, 200 to 3,000 partially diffused designs were generated.

The new backbones were then subjected to ProteinMPNN sequence design and AF2 predictions, as previously described. These designs were filtered using the same criteria, with the observation that partial RFDiffusion tended to increase the overall number of passing designs. However, it was also noted that the original hydrogen bonding networks (particularly bidentate hydrogen bonds to the target backbone) were sometimes disrupted without being detected by AF2. To address this, we implemented an additional filter, ‘buns’ (buried unsatisfied hydrogen bonds) <=1, to explicitly count unsatisfied heavy atoms. Depending on the target’s polarity, this step could filter out 50-80% of designs.

#### Parametric perturbation

Helices that were suboptimal in terms of interaction with the target could be isolated from the binder template and underwent defined x/y/z translations and rotations. Rosetta-based scoring of these translations and rotations filtered perturbations that were either non-productive (moved the helix too far away from the target) or clashed with the target. Then motif RFDiffusion was used to connect the perturbed helix to the rest of the binder template.

### Peptide generation

All Fmoc-protected amino acids were obtained from P3 Biosystems; the coupling reagent HATU, DIPEA, and other chemicals unless otherwise stated were obtained from Sigma-Aldrich; DIC was obtained from Oakwood Chemicals; acetonitrile (ACN), methylene chloride (DCM), diethyl ether, and DMF were obtained from Fisher Scientific. Preloaded Wang resin and OxymaPure were obtained from CEM.

The linear peptide sequences were synthesized via solid-phase peptide synthesis (SPPS) on a LibertyBlue microwave synthesizer (CEM) at 0.1mmol scale on preloaded Wang resin. The complete linear peptide sequence was then coupled by hand to Fmoc-6-aminohexanoic acid (AHX) (3eq), HATU (3 eq), and DIPEA (5eq) for 3h, washed with DMF, then treated with 20% piperidine in DMF for 2×15m to remove Fmoc. The resin was then washed with DMF and swelled in 50/50 DMSO/DMF prior to coupling with biotin (3eq), HATU (eq), and DIPEA (5eq) for 3h, then washed with DMF and DCM prior to total deprotection and cleavage from resin using a cleavage cocktail (92.5:2.5:2.5:2.5 TFA:triisopropylsilane:H_2_O:2,2’-(ethylenedioxy)diethanethiol, v/v/v/v) for 3h. The peptide cleavage solution was concentrated *in vacuo* and precipitated in ice-cold diethyl ether, centrifuged to pellet the peptide crude (7000g, 4°C, 10m), and dried under N_2_. The peptide crude was resuspended in minimal water and acetonitrile and purified by RP-HPLC on a semi-prep Agilent 1260 Infinity with a linear gradient from 10-45% A->B (A: water with 0.1% TFA, B: ACN with 0.1% TFA) on a Zorbax C18-300SB 5 μm 9.4×250mm column (Agilent), and analyzed via LC/MS-TOF on an Agilent G6230B. Pure peptide fractions were combined and lyophilized.

### DNA library preparation

All protein sequences were padded to a uniform length by adding a (GGGS)n linker at the C terminal of the designs, to avoid the biased amplification of short DNA fragments during PCR reactions. The protein sequences were reversed translated and optimized using DNAworks2.0 with the S. cerevisiae codon frequency table. Homologous to the pETCON plasmid Oligo libraries encoding the designs were ordered from Twist Bioscience. Combinatorial libraries were ordered as IDT (Integrated DNA Technologies) ultramers with the final DNA diversity ranging from 1×10^6^ to 1×10^7^.

All libraries were amplified using Kapa HiFi Polymerase (Kapa Biosystems) with a qPCR machine (BioRAD CFX96). In detail, the libraries were firstly amplified in a 25 μL reaction, and PCR reaction was terminated when the reaction reached half the maximum yield to avoid over-amplification. The PCR product was loaded to a DNA agarose gel. The band with the expected size was cut out and DNA fragments were extracted using QIAquick kits (Qiagen, Inc.). Then, the DNA product was re-amplified as before to generate enough DNA for yeast transformation. The final PCR product was cleaned up with a QIAquick Clean up kit (Qiagen, Inc.). For the yeast transformation, 2-3 μg of digested modified pETcon vector (pETcon3) and 6 μg of insert were transformed into EBY100 yeast strain using the protocol as described before.

DNA libraries for deep sequencing were prepared using the same PCR protocol, except the first step started from yeast plasmid prepared from 5×10^7^ to 1×10^8^ cells by Zymoprep (Zymo Research). Illumina adapters and 6-bp pool-specific barcodes were added in the second qPCR step. Gel extraction was used to get the final DNA product for sequencing. All libraries include the native library and different sorting pools were sequenced using Illumina NextSeq/MiSeq sequencing.

For mammalian cell expression, all plasmids use the pcDNA3.1 backbone and were produced by GenScript.

### Yeast surface display

*S. cerevisiae* EBY100 strain cultures were grown in C-Trp-Ura media and induced in SGCAA media following the protocol as described before. Cells were washed with PBSF (PBS with 1% BSA) and labeled with biotinylated target using two labeling methods, with-avidity and without-avidity labeling. For the with-avidity method, the cells were incubated with biotinylated target, together with anti-c-Myc fluorescein isothiocyanate (FITC, Miltenyi Biotech) and streptavidin–phycoerythrin (SAPE, ThermoFisher). The concentration of SAPE in the with-avidity method was used at 1/4 concentration of the biotinylated target. The with-avidity method was used in the first few rounds of screening of the original design to fish out weak binder candidates. For the without-avidity method, the cells were firstly incubated with biotinylated target, washed, secondarily labeled with SAPE and FITC. For these designs, the first two to four rounds of sorts were applied with 1 μM concentration of the Ras C-terminus with biotin at the C-terminal end of the peptide. The remaining subsequent sorts were done with varying concentrations (1 nM - 1 μM) of full length RAS. The final sorting pools of the combinatorial libraries were sequenced using Illumina NextSeq/MiSeq sequencing. All FACS data was analyzed in FlowJo.

### Cell culture and transfection

HEK293T, HeLa, MEF, Rasless MEF (isogenic cell lines)^19^, MIA PaCa-2, H358, H441, Rh36, RD, H1299 cell lines were cultured in Dulbecco’s modified Eagle medium (DMEM) containing 1 g L^-1^ glucose and supplemented with 10% (v/v) fetal bovine serum (FBS) and 1% (v/v) penicillin–streptomycin (Pen-Strep). H441, H2228, and MOLM-13 cells were grown in RPMI-1640 cell media containing 1 g L^-1^ glucose and supplemented with 10% (v/v) fetal bovine serum (FBS) and 1% (v/v) penicillin–streptomycin (Pen-Strep). For MOLM-13 cells, 2mM glutamine was also added to the media. All cells were grown in a humidified incubator at 5% CO_2_ and at 37°C.

Before transfection, all cells were plated onto sterile poly-D-lysine coated plates or dishes and grown to 50%–70% confluence. All cells were transfected with Lipofectamine LTX. Subsequently, all cells were grown for an additional 1-2 days before experimental testing. All cells underwent serum starvation for 16 hours before downstream assay analysis. See **Table S2** for details of reagents.

### General procedures for bacterial protein production and purification

The *E. coli* Lemo21(DE3) strain was transformed with a pET29b^+^ plasmid encoding the synthesized gene of interest. Cells were grown for 24 hours in liquid broth medium supplemented with kanamycin. Cells were inoculated at a 1:50 ml ratio in the Studier TBM-5052 autoinduction medium supplemented with kanamycin, grown at 37°C for 2–4 hours and then grown at 18°C for an additional 18 hours. Cells were collected by centrifugation at 4,000 *g* at 4 °C for 15 min and resuspended in 30 ml lysis buffer (20 mM Tris-HCl, pH 8.0, 300 mM NaCl, 30 mM imidazole, 1 mM PMSF and 0.02 mg ml^-1^ DNase). Cell resuspensions were lysed by sonication for 2.5 min (5 s cycles). Lysates were clarified by centrifugation at 24,000 *g* at 4°C for 20 min and passed through 2-ml Ni-NTA nickel resin pre-equilibrated with wash buffer (20 mM Tris-HCl, pH 8.0, 300 mM NaCl and 30 mM imidazole). The resin was washed twice with 10 column volumes (Cversus) of wash buffer, and then eluted with 3 Cversus elution buffer (20 mM Tris-HCl, pH 8.0, 300 mM NaCl and 300 mM imidazole). The eluted proteins were concentrated using Ultra-15 Centrifugal Filter Units and further purified by using a Superdex 75 Increase 10/300 GL size exclusion column in TBS (25 mM Tris-HCl, pH 8.0, and 150 mM NaCl). Fractions containing monomeric protein were pooled, concentrated and snap-frozen in liquid nitrogen and stored at −80°C. See **Table S2** for details of reagents.

To biotinylate RIBs, an AviTag was added at their C-terminus. Biotinylation of purified RIBs fused with AviTags was performed using the BirA bulk kit according to manufacturer’s protocol (Avidity LLC). Briefly, biotinylation reactions (pH 8.0; 1:1 ratio) were performed for 1 hour at 4°C on an orbital shaker and then excess biotinylation reagent was removed using Superdex 200 Increase 10/300 GL (depending on kDa of protein) size exclusion column in TBS (25 mM Tris-HCl, pH 8.0, and 150 mM NaCl).

### Biolayer interferometry

Protein–protein interactions were measured by using an Octet RED96 System (ForteBio) using streptavidin-coated biosensors (ForteBio). Each well contained 200 μL of solution, and the assay buffer was HBS-EP+ buffer (GE Healthcare Life Sciences, 10 mM HEPES pH 7.4, 150 mM NaCl, 3 mM EDTA, 0.05% (v/v) surfactant P20) plus 0.5% non-fat dry milk blotting grade blocker (BioRad). The biosensor tips were loaded with analyte peptide or protein at 20 μg mL^-1^ for 300 s (threshold of 0.8 nm response), incubated in HBS-EP+ buffer for 60 s to acquire the baseline measurement, dipped into the solution containing cage and/or key for 1800 s (association step) and dipped into the HBS-EP+ buffer for 1800 s (dissociation steps). The binding data were analyzed with the ForteBio Data Analysis Software version 9.0.0.10.

### Immunostaining

All cell lines were seeded onto 24-well glass-bottom plates. After transfection and drug addition, cells were fixed with 4% PFA in 2x PHEM buffer (60 mM PIPES, 50 mM HEPES, 20 mM EGTA, 4 mM MgCl_2_, 0.25 M sucrose, pH 7.3) for 10 min, permeabilized with 100% methanol for 10 min, washed with PBS 3x, blocked in 1% BSA in PBS for 30 min, incubated with biotinylated RIBs and antibodies for 2 hours at room temperature, washed with PBS 3x, incubated with DAPI and fluorescently labeled reagents (streptavidin and BioTracker) for 1 hour at room temperature and aluminum foil cover. Cells were then washed with PBS 3x and mounted for epifluorescence imaging. All images were analyzed in ImageJ. See **Table S2** for details of reagents.

### Immunoblotting and Immunoprecipitation

Cells expressing indicated constructs and incubated with indicated drugs were plated, transfected, and labeled as described in figure legends. For cells that required membrane fractionation were treated with the Cell Fractionation Kit from Cell Signaling Technologies according to the manufacturer’s protocol. Cells were then transferred to ice and washed 2x with ice cold DPBS. Cells were then detached from the well by addition of 1x RIPA lysis buffer (50 mM Tris pH 8, 150 mM NaCl, 0.1% SDS, 0.5% sodium deoxycholate, 1% Triton X-100, 1x protease inhibitor cocktail, 1 mM PMSF, 1mM Na_3_VO_4_, 1% NP-40) and either scraping of cells or rotation on shaker for 30 min at 4°C. Cells were then collected and vortexed for at least 5 s every 10 min for 20 min at 4°C. Cells were then collected and clarified by centrifugation at 13,000g for 10 minutes at 4°C. The supernatant was collected and underwent Pierce BCA assay to quantify total protein amounts.

For immunoblotting, whole cell lysate protein amounts were normalized across samples in the same gel, mixed with 4x loading buffer prior to loading, incubated at 95°C for 5 min and then 4°C for 5 min, and separated on Any kDa SDS-PAGE gels. Proteins separated on SDS-page gels were transferred to nitrocellulose membranes via the TransBlot system (BioRad). The blots were then blocked in 5% milk (w/v) in TBST (Tris-buffered saline, 0.1% Tween 20) for 1 hour at room temperature. Blots were washed with TBST 3x then incubated with indicated primary antibodies or biotinylated RIBs in 1% BSA (w/v) in TBST overnight at 4°C. Blots were then washed with TBST 3x and incubated with LICOR dye-conjugated secondary antibodies (LICOR 680/800 or streptavidin-LICOR 800) and fluorescently labeled streptavidin in 1% BSA (w/v) in TBST for 1 hour at room temperature. The blots were washed with TBST 3x and imaged on an Odyssey IR imager (LICOR). Quantitation of Western blots was performed using ImageJ on raw images. See **Table S2** for details of reagents including antibody dilutions.

For immunoprecipitation, streptavidin beads were loaded by three lysis buffer washes before the addition of 1 mg ml^−1^ of biotinylated RIBs at 4 °C on an orbital shaker for 3 h. Beads were then washed two times in lysis buffer. Whole-cell lysate protein amounts were normalized across samples and protein samples were added to beads (at least 100 μg per sample) either at room temperature for 1 h for streptavidin beads or at 4 °C on an orbital shaker overnight. Beads were then washed two times in lysis buffer and one time in TBS and then mixed with 4× loading buffer. The remaining portion of the protocol was the same as for immunoblotting.

### Epifluorescence imaging

Cells were washed twice with FluoroBrite DMEM imaging media and subsequently imaged in the same media in the dark at room temperature. Epifluorescence imaging was performed on a Yokogawa CSU-X1 spinning dish confocal microscope with either a Lumencor Celesta light engine with 7 laser lines (408, 445, 473, 518, 545, 635, 750 nm) or a Nikon LUN-F XL laser launch with 4 solid state lasers (405, 488, 561, 640 nm), 40x/0.95 NA objective or 60x/1.4 NA oil immersion objective and a Hamamatsu ORCA-Fusion scientific CMOS camera, both controlled by NIS Elements 5.30 software (Nikon). The following laser and filter combinations (center/bandwidth in nm) were used: GFP: EX473 EM525/36, RFP: EX545 EM605/52, BFP: EX445 EM525/36. Exposure times were 500ms for all channels, with no EM gain set and no ND filter added. All epifluorescence experiments were subsequently analyzed using Image J. Brightfield images were acquired on the ZOE Fluorescent Cell Imager (BioRad). See **Table S2** for details of reagents.

### Co-localization analysis

For co-localization analysis, cell images were individually thresholded and underwent Coloc 2 analysis on ImageJ. Mander’s coefficient, which ranges from 0 to 1 with 1 being 100% colocalized, is measuring the spatial overlap of one imaging channel with another imaging channel. Pearson’s coefficient compares the pixel intensity of one channel with another channel. Pearson’s coefficient values can range from −1 to 1 with −1 meaning inversely proportional and 1 meaning same pixel intensities.

### MOLM-13 cell line generation and cell growth assays

MOLM-13 KRAS and NRAS mutant cell lines were generated as previously described^29,30^. Briefly, the plasmids pCW57.1 (Addgene 41393), pDONR223 KRAS WT (Addgene 81751), and pDONR223 NRAS WT (Addgene 82151) were used to generate doxycycline-inducible KRAS and NRAS constructs. mCherry was cloned (Addgene 60954) to the N-terminus of KRAS or NRAS on the pCW57.1 backbone and site-directed mutagenesis was performed to generate KRAS and NRAS mutations. MOLM-13 cells (DSMZ) were transduced with lentivirus generated from these constructs and sorted for mCherry positivity after treatment with doxycycline 2 μg/mL. MOLM-13 cells were transfected with plasmids and treated with doxycycline 2 µg/mL (if indicated). 6 hours later, 10 nM AC220 (quizartinib) was added and this addition marks the start of the cell counting (t=0 days). Cell counts were obtained using a Countess 3 automated cell counter (ThermoFisher).

### Graphics

All schematics were generated using BioRender.

### Statistics and reproducibility

No statistical methods were used to predetermine the sample size. No sample was excluded from data analysis, and no blinding was used. All data were assessed for normality. For normally distributed data, pairwise comparisons were performed using unpaired two-tailed Student’s t tests, with Welch’s correction for unequal variances used as indicated. Comparisons between three or more groups were performed using ordinary one-way or two-way analysis of variance (ANOVA) as indicated. For data that were not normally distributed, pairwise comparisons were performed using the Mann-Whitney U test, and comparisons between multiple groups were performed using the Kruskal-Wallis test. All data shown are reported as mean ± SEM and error bars in figures represent SEM of biological triplicates. All data were analyzed and plotted using GraphPad Prism 8 including non-linear regression fitting.

### Data availability

The data that support the findings of this study will be available from Figshare.

### Code availability

The code and design models used in this study will be available from Zenodo.

## Supplemental Figures

**Figure S1:**
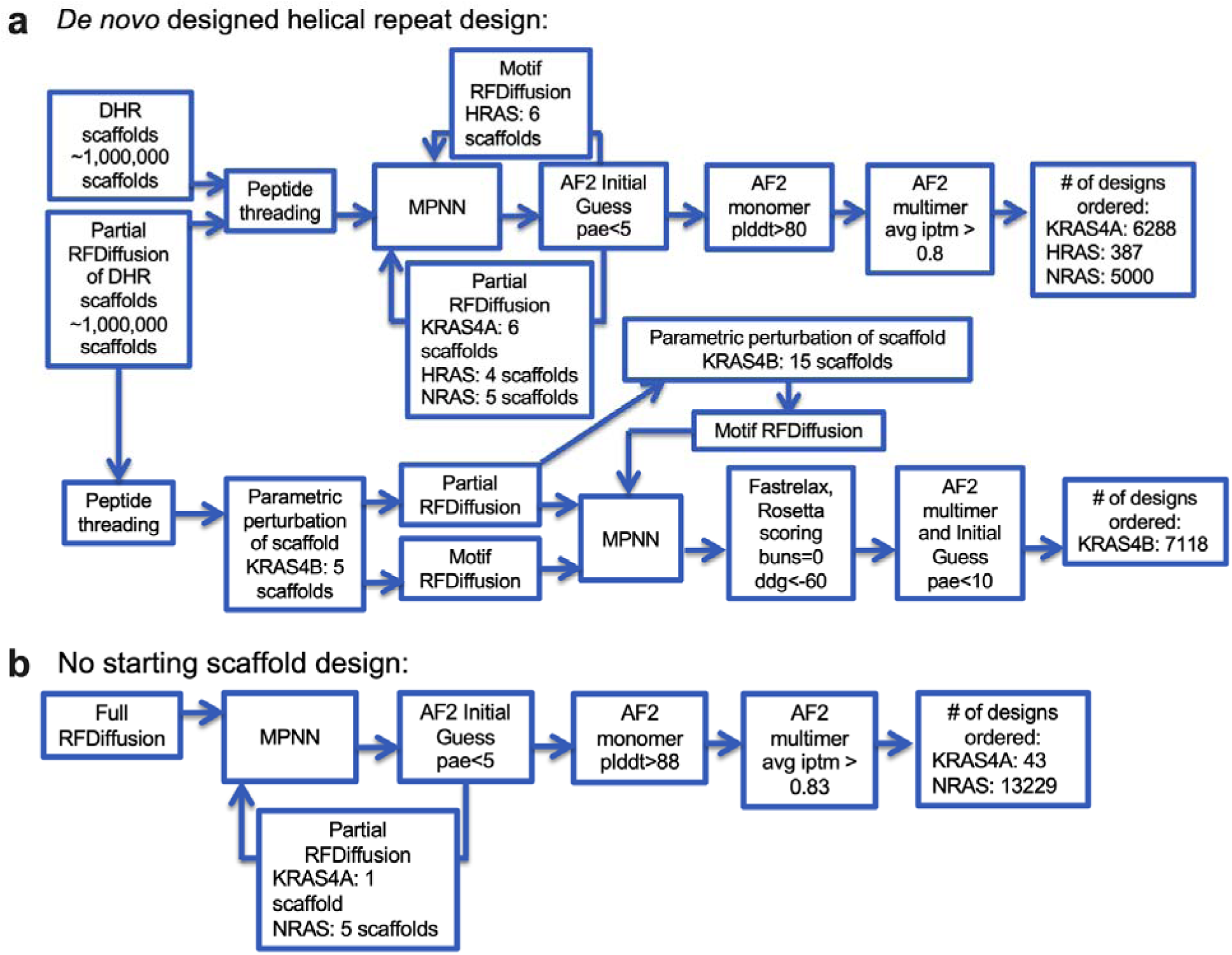
Details of computational design pipeline. **a-b**) Details of the computational design workflow for designing RIBs either using (**a**) *de novo* designed helical repeat proteins for the logos pipeline as starting scaffolds. (**b**) RIBs were also designed without a particular starting scaffold utilizing full two-sided RFDiffusion with both strand and helix specification.

**Figure S2:**
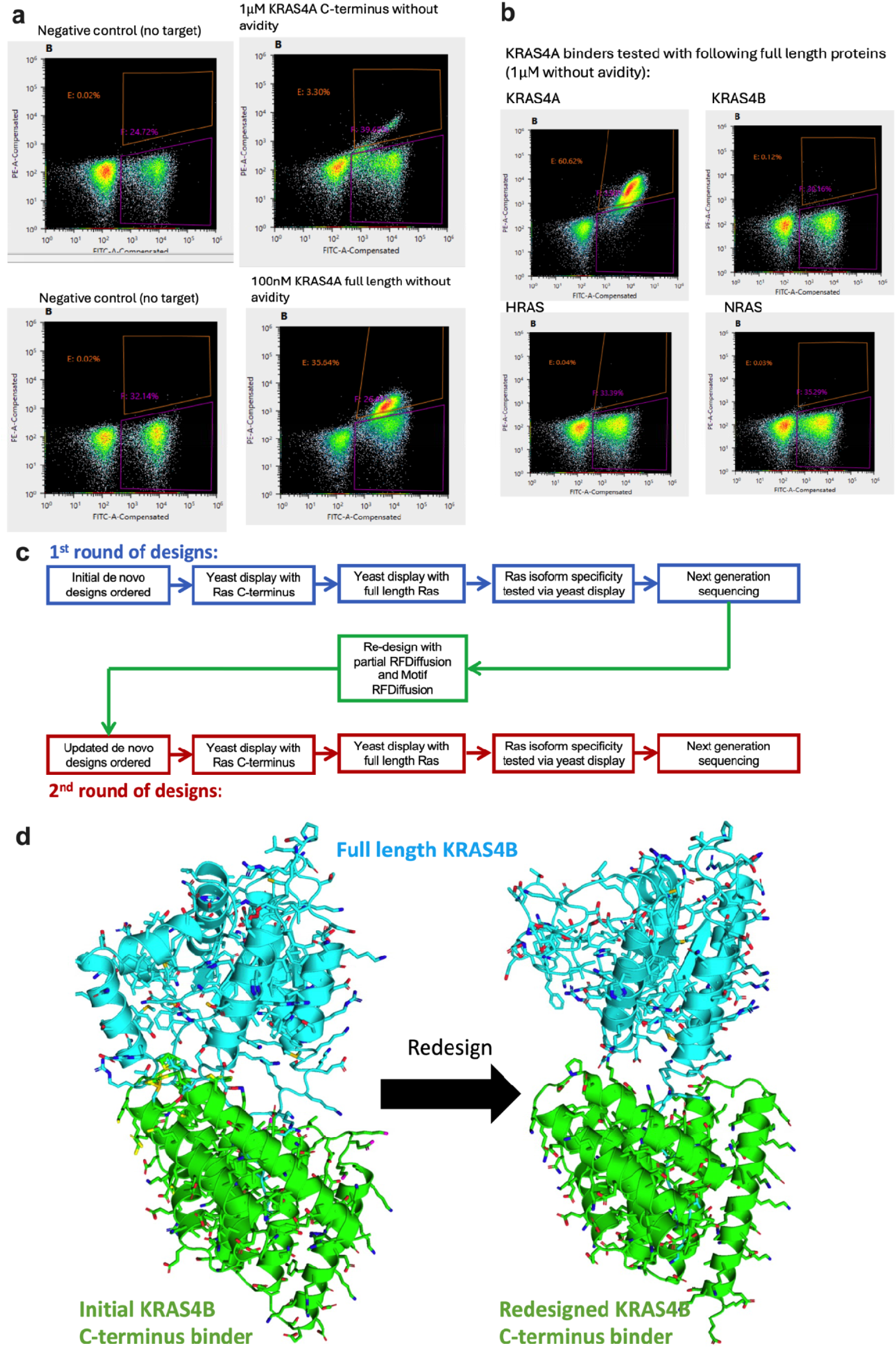
High throughput yeast display testing of designs. **a-b**) Representative flow cytometry plots of designs tested via yeast display (see Methods for details). X-axis represents binder display and Y-axis represents target binding. Negative controls are when no target was added. (**a**) Either biotinylated C-terminus or full length protein was incubated with yeast cells expressing on their surface the designs. (**b**) Testing of Ras isoform specificity was initially done by incubating biotinylated full length Ras isoforms with yeast display by incubating yeast cells expressing designs. For all cases, none of the design libraries had significant binding signal to off-target Ras isoforms. **c)** Experimental and computational workflow for designing RIBs. The first round of computational design focused only on the Ras C-terminus and were tested for binding to Ras C-terminus and full length protein via yeast display. For the RIBs that only bound to the C-terminus (KRAS4B and NRAS), the binder backbones were optimized via partial RFDiffusion and motif RFDiffusion for both the C-terminus (to tighten target binding) and the full length protein (to account for steric clashes). After this second round of computational design, these designs were again tested for binding to Ras C-terminus and full length protein via yeast display. **d)** AlphaFold2 structure predictions of KRAS4B RIBs. Left: KRAS4B C-terminus binder which is predicted to sterically clash with the folded domain of KRAS4B. Right: After redesign of these KRAS4B C-terminus binder, a new KRAS4B RIB design is predicted to no longer clash with full length KRAS4B.

**Figure S3:**
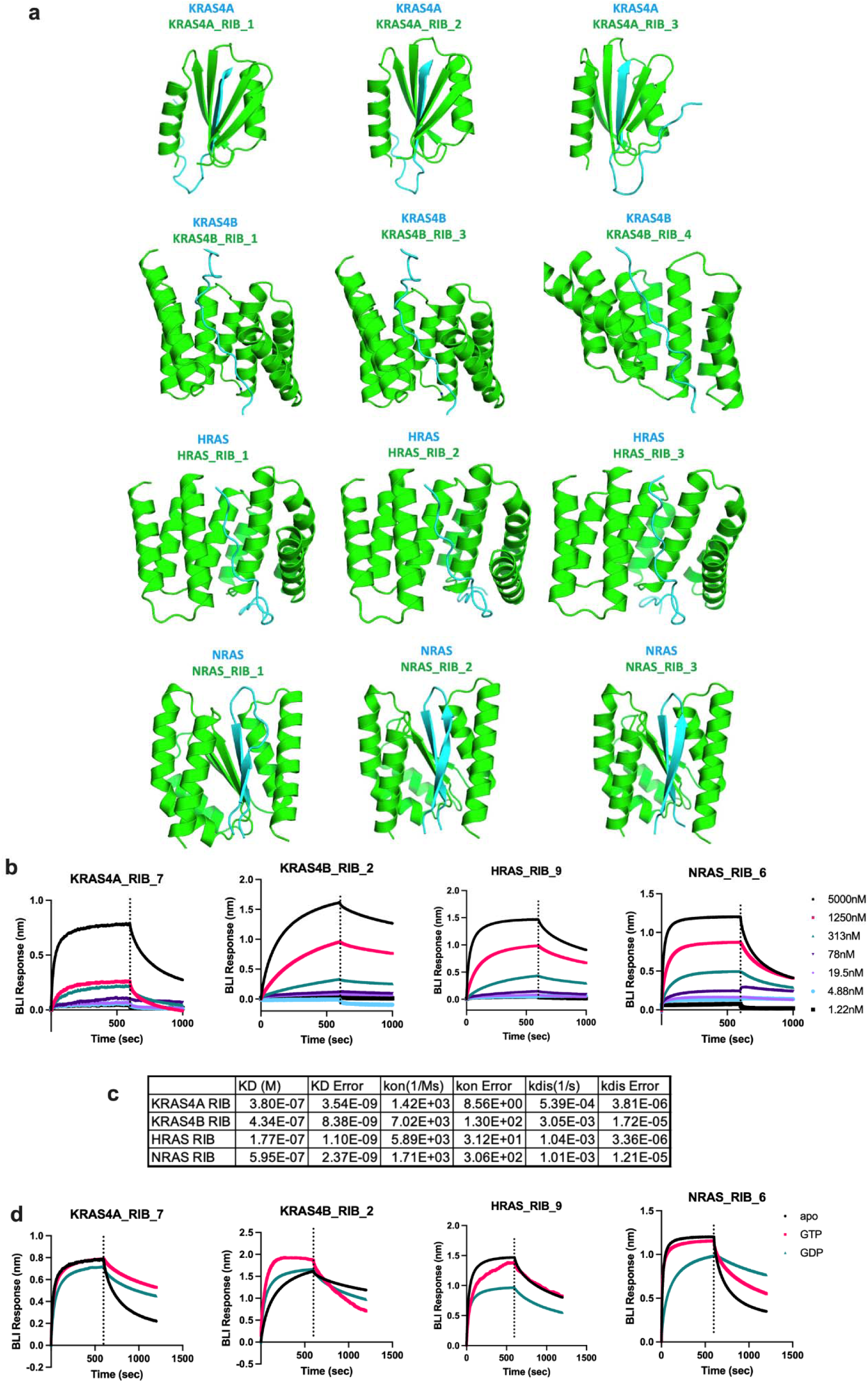
*In vitro* binding of RIBs to their target. **a)** Design models of representative RIBs tested for each target. **b)** BLI results of RIBs binding to their full length target. Results are representative of 3 independent experiments. The BLI data is for the RIB that was used for the rest of the paper. **c)** Estimated binding kinetics and affinity of RIBs with their full length target. **d)** BLI results of RIBs (5 M) binding to their full length target either without nucleotide (apo), 5 M GTP (GTP), or 5 M GDP (GDP). Results are representative of 3 independent experiments.

**Figure S4:**
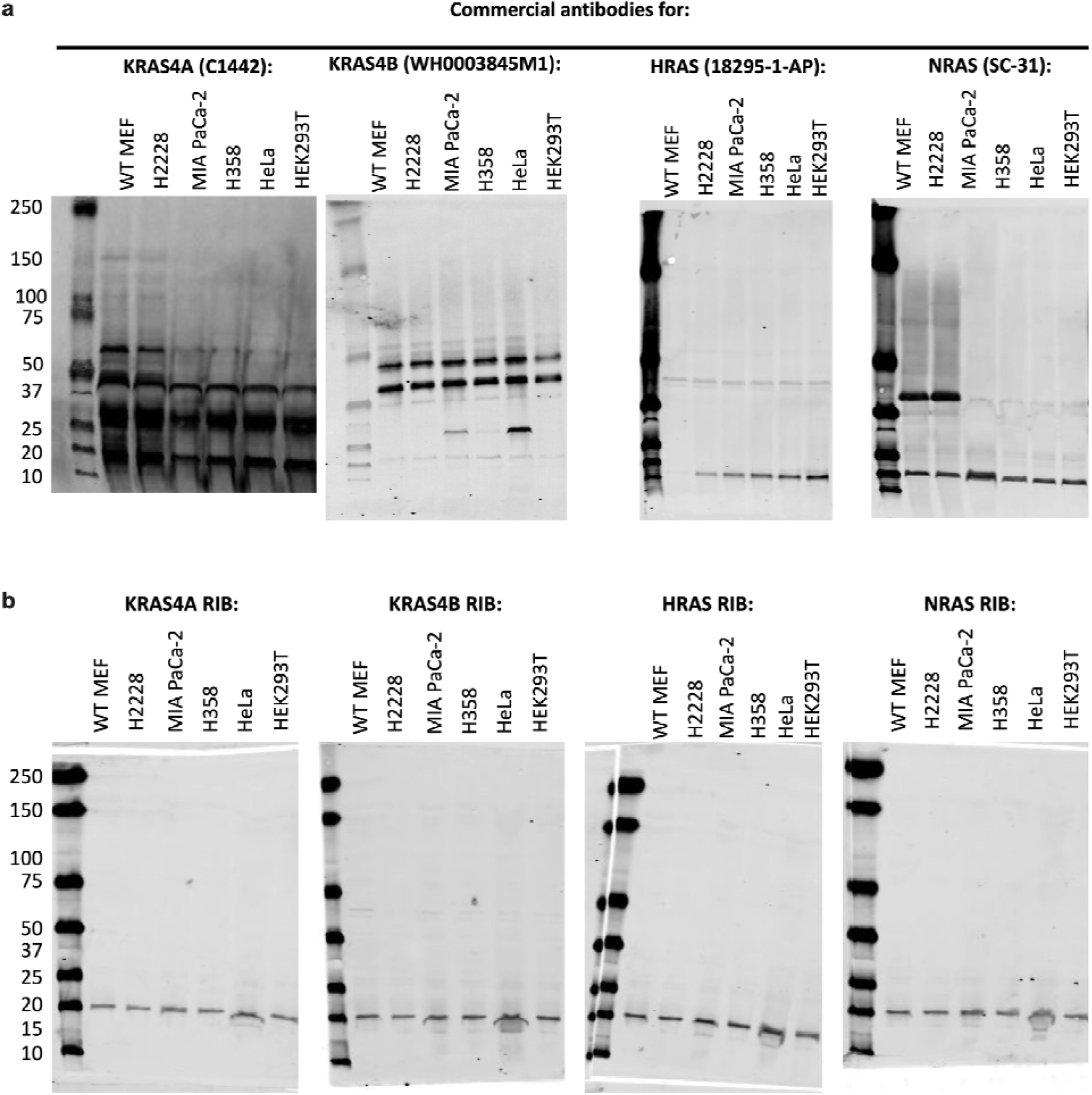
Comparison to commercially available antibodies. **a)** Lysates from a panel of cell lines were lysed, ran on SDS-PAGE gels, and probed with antibodies that were shown previously to be the most specific among the commercially available reagents. Shown are representative full blots from 3 independent experiments. **b)** Lysates from a panel of cell lines were ran on SDS-PAGE gels and probed with biotinylated RIBs. Shown are representative full blots from 3 independent experiments.

**Figure S5:**
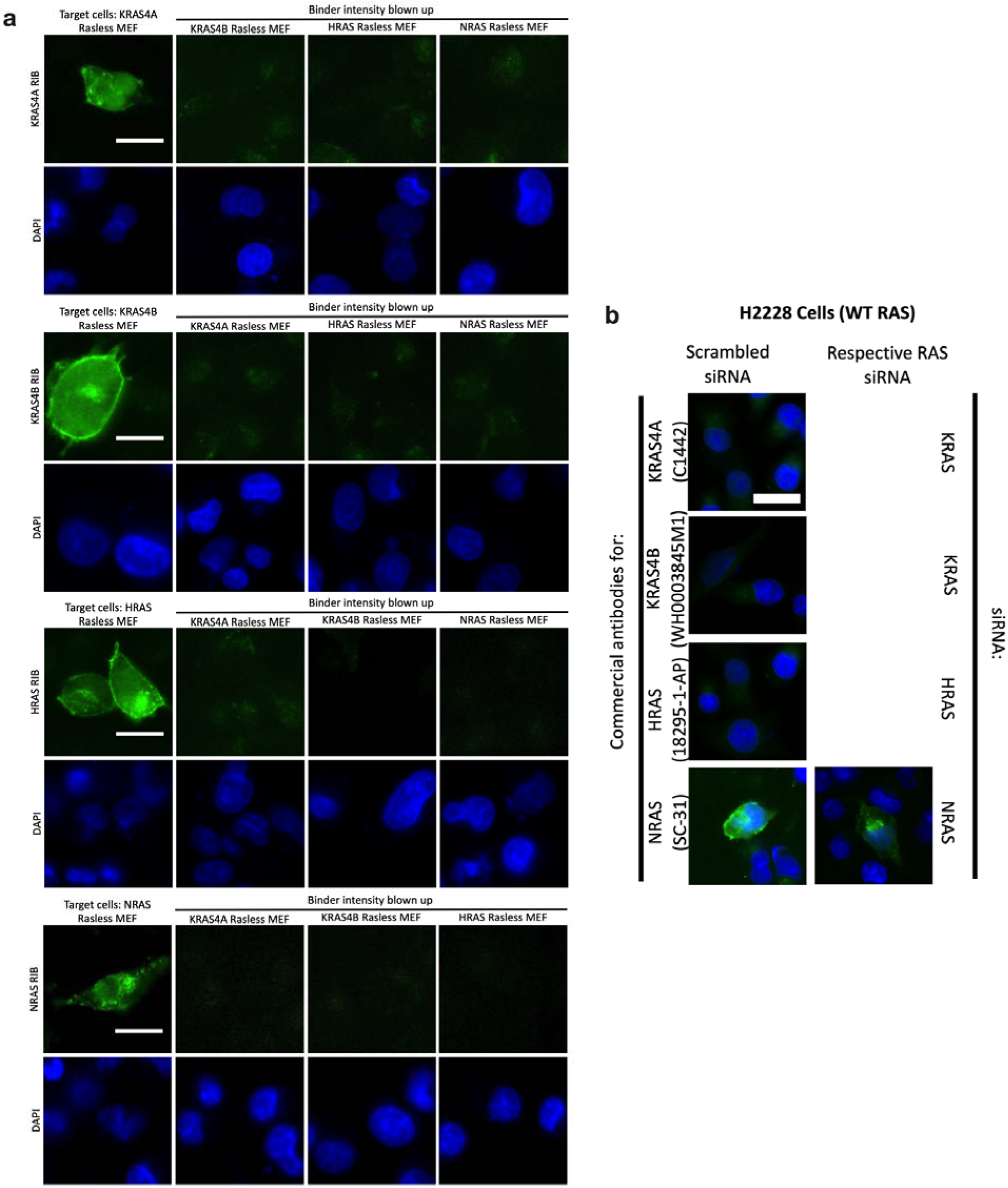
RIBs specifically bind to their target in mammalian cells. **a)** Rasless MEFs were fixed, permeabilized, and probed with biotinylated RIBs and DAPI. Shown are the separated epifluorescence signals from RIBs and DAPI. RIBs incubated with their target cells are shown with a fixed brightness and contrast while RIBs incubated with their off-target cells are shown with the background signal amplified. Images are the same as Fig 3f. Results are representative of 3 independent experiments. Scale bar=10 m. **b)** Representative epifluorescence images of Rasless MEFs either transfected with scrambled or on target siRNAs for 2 days and then immunostained with antibodies that were shown previously to be the most specific among the commercially available reagents. Results are representative of 3 independent experiments. Scale bar=10µm.

**Figure S6:**
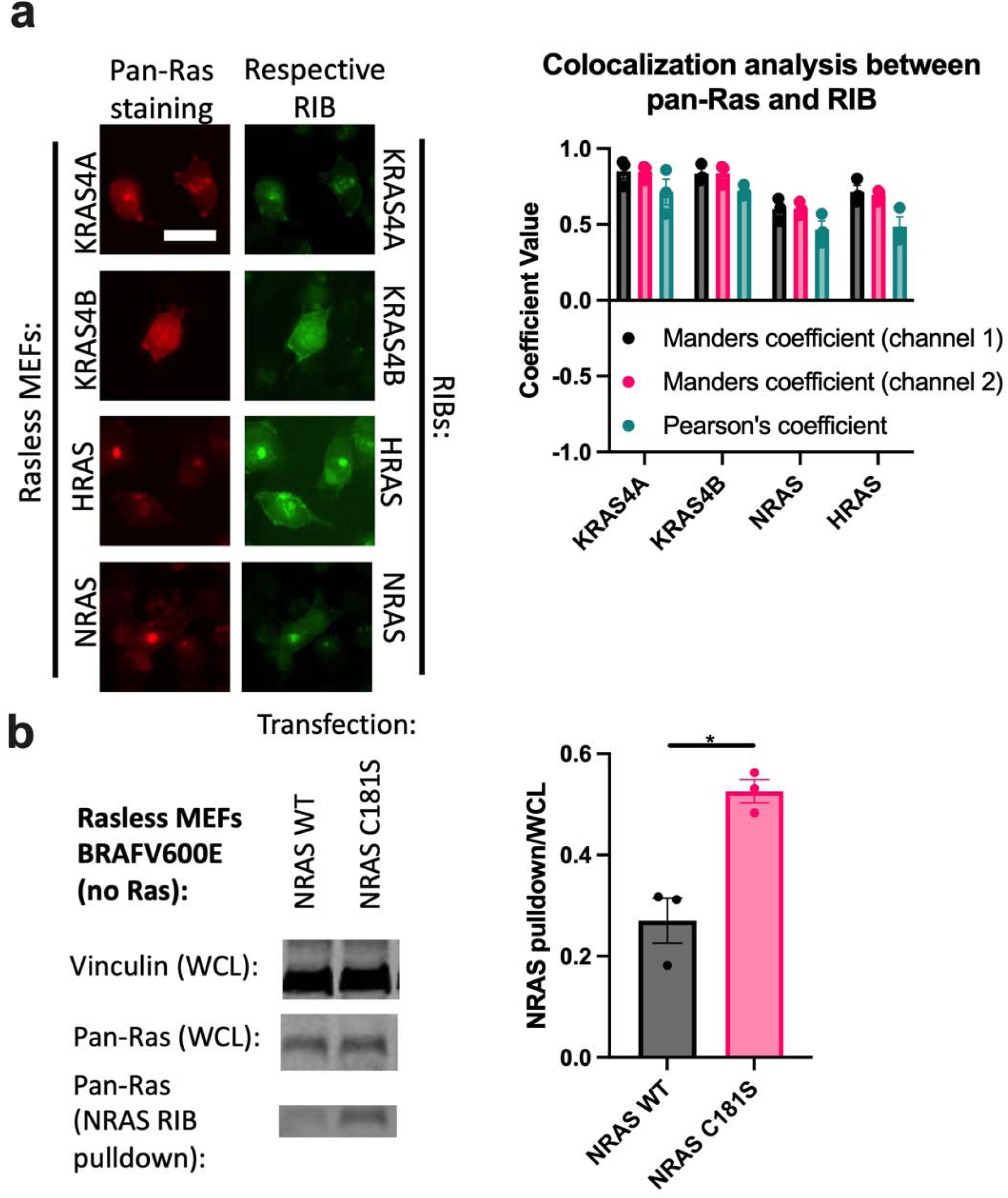
RIBs evenly bind to target throughout the mammalian cell. **a)** Rasless MEFs were fixed, permeabilized, probed via immunostaining with the respective biotinylated RIBs and pan-Ras, and then imaged. Images were analyzed for co-localization between pan-Ras and RIB signal (n=3 experiments). Images are representative of 3 independent experiments. Scale bar=10µm. **b)** Rasless MEFs harboring BRAFV600E express none of the major Ras isoforms. These cells were transfected with either NRAS WT or NRAS C181S for 1 day, lysed, either pulled down with beads loaded with NRAS RIBs (see Methods for details) or had no pulldown (whole cell lysates=WCL), and then immunoprobed with the indicated antibodies. Left: representative immunoblot. Right: densitometry quantification of immunoblots (n=3 experiments). Statistics: Unpaired 2-tailed student’s t-test.

**Figure S7:**
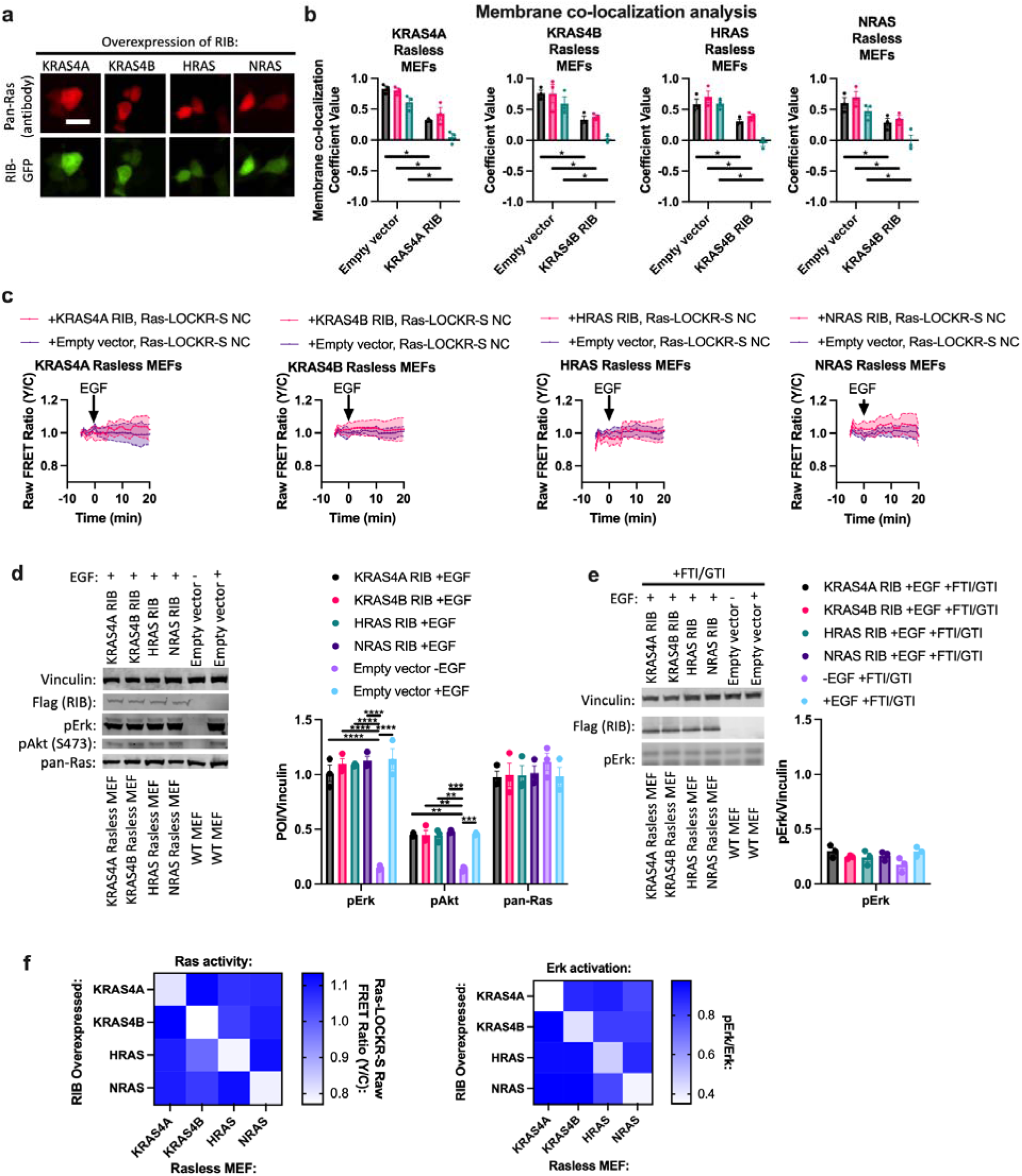
RIBs affect Ras signaling. **a)** Epifluorescence images of Rasless MEFs expressing RIB fused to GFP (RIB-GFP). These cells were then immunostained for pan-Ras. Images are representative of three biologically independent experiments. Scale bar=10 m. **b)** Rasless MEFs expressing RIB-GFP were stained for pan-Ras and BioTracker Cytosolic Membrane dyes. Co-localization between RIB-GFP signal and BioTracker signal was analyzed (n=3 experiments). Statistics: Unpaired 2-tailed student’s t-test. **c)** Time course imaging of Ras-LOCKR-S negative control (NC) expressed in Rasless MEFs either co-expressing a RIB or empty vector and stimulated with 100ng/mL EGF (n=10 cells for each experiment). These curves are representative of three biologically independent experiments. **d-e**) Rasless MEFs were transfected with their respective RIB and either treated without (**d**) or with (**e**) 10µM dual farnesyl transferase inhibitor (FTI) and geranylgeranyl transferase-1 inhibitor (GTI) FGTI-2734^31^. 1 day later, cells were then stimulated with 100ng/mL EGF. These cells were compared to WT MEFs transfected with empty vector and stimulated with 100ng/mL EGF. These cells then underwent immunoblotting analysis. Right: densitometry quantification of immunoblots (n=3 experiments). Statistics: two-way ordinary ANOVA. **f**) Rasless MEFs were transfected with RIBs for 1 day. Left: To measure Ras activity, Ras-LOCKR-S was co-transfected and imaged the next day. Right: To measure pErk levels, cells underwent immunoblotting analysis and pErk levels were compared to total Erk levels. All experiments were done in triplicates.

**Figure S8:**
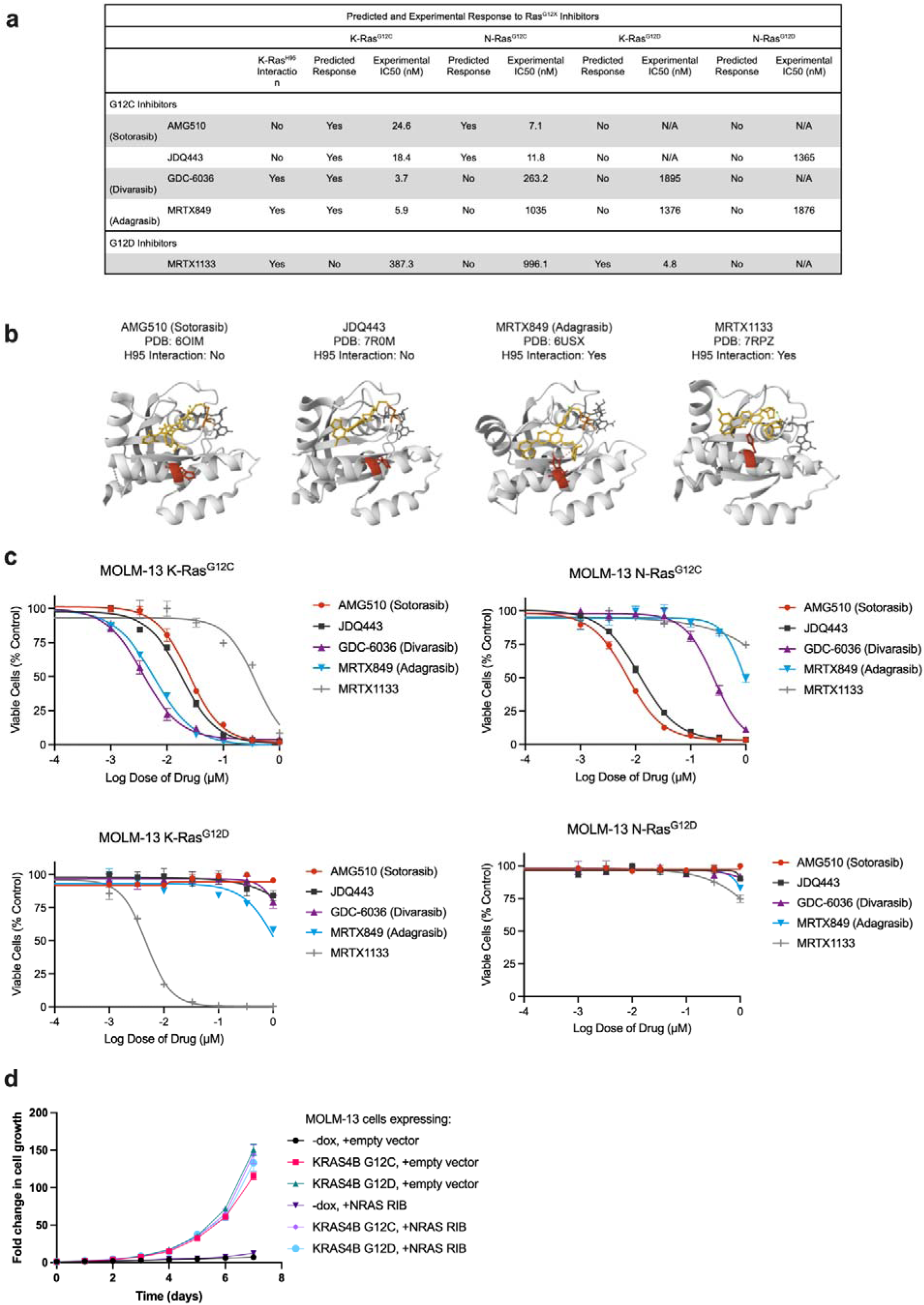
Characterization of MOLM-13 cells. **a)** As previously described^32,33^, the selectively of K-Ras^G12C^ and K-Ras^G12D^ inhibitors over N-Ras mutant counterparts depends upon whether a drug binds to K-Ras H95, a residue that is not conserved in N-Ras (L95). Experimental IC50s are based on (**c**). **b)** Published crystal structures of K-Ras bound to G12C and G12D inhibitors showing the position of the drug in relation to H95 (in red). The structure for K-Ras bound to GDC-6036 (Divarasib) has not yet been published, but molecular dynamic simulations suggest that GDC-6036 binds to H95 (https://chemrxiv.org/engage/chemrxiv/article-details/64fecb59b6ab98a41c3d9c0f). **c)** MOLM-13 KRAS or NRAS mutant cell lines treated with K-Ras^G12C^ or K-Ras^G12D^ inhibitors. Represented results shown from 3 independent experiments each performed in technical triplicate. **d)** MOLM-13 cell lines expressing either KRAS4B G12C or G12D were treated with or without doxycycline to turn on mutant KRAS4B expression. Cells were also transfected with NRAS RIB expressing plasmid or empty vector and co-treated with AC220. Cells were then counted over a 7 day period (n=3 experiments). Statistics: one-way ordinary ANOVA.

**Table S1: Characterization of all RIBs experimentally tested *in vitro* and in cells**

The top 10 RIBs from yeast display for each target were further tested *in vitro* for binding affinity and in cells for their effects on Ras expression (via immunoblot), Ras activity (via Ras-LOCKR-S) before and after 100ng/mL EGF stimulation, pErk signaling (via immunoblot) before and after 100ng/mL EGF stimulation, localization of signal when used as staining probes, and number of bands when applied in blots.

**Table S2: Reagents used within study**

Reagents used throughout the study with catalog numbers.

